# Layer-specific glutamatergic inputs and Parvalbumin interneurons modulate early life stress induced alterations in prefrontal glutamate release during fear conditioning in pre-adolescent rats

**DOI:** 10.1101/2025.09.08.674892

**Authors:** Jiamin Song, Muzammil Younus, Hong Long, Tak Pan Wong, Claire-Dominique Walker

## Abstract

Exposure to early life stress (ELS) can exert long-lasting impacts on emotional regulation. The corticolimbic system including the basolateral amygdala (BLA), ventral hippocampus (vHIP), and the medial prefrontal cortex (mPFC) plays a key role in fear learning. Using the limited bedding paradigm (LB), we examined the functional consequences of ELS on excitatory and inhibitory tone in the prelimbic (PL) mPFC after fear conditioning in rats. In adults, LB exposure enhanced *in vivo* glutamate release in the PL mPFC during fear conditioning in male, but not female offspring. In contrast, the glutamate response to fear conditioning was diminished in LB-exposed pre-adolescent males, but not females. We investigated whether reduced glutamatergic inputs and/or elevated inhibitory tone might contribute to the diminished glutamate response in the mPFC following LB in pre-adolescent male rats. Indeed, we found that LB exposure specifically increased the activation of PV, but not SST interneurons in layer V, but not layer II/III of the PL mPFC in fear-exposed pre-adolescent males. Presynaptic glutamate release probability was reduced by LB exposure in layer V, but increased in layer II/III of the PL mPFC. These functional changes might be related to the LB-induced alterations in the bilaminar distribution of BLA and vHIP projections to the PL mPFC we observed in pre-adolescent males. Overall, our findings suggest that ELS modifies glutamate release and PL mPFC function during fear conditioning in a sex- and age-dependent fashion, likely through layer-specific shifts in excitation/inhibition balance.

**Significance Statement:** Early life stress (ELS) increases the risk of developing affective disorders and long-term emotional dysregulation might arise from disruptions in the development of the fear circuitry. This study examines how ELS modifies fear-induced activity of long-range excitatory projections and local inhibitory microcircuits in the developing prefrontal cortex. We tested whether ELS-induced alterations in prefrontal cortex function are sex- and age-dependent, leading to the well-documented sex differences in emotional behavioral outcome. Studying how ELS differentially modifies regional excitatory inputs and cell type specific activation in the prefrontal cortex during a critical period of brain development will enhance our understanding of the neurobiological mechanisms underlying the pathogenesis of emotional dysregulation and inspire more targeted intervention after exposure to early adversity.

## Introduction

Early life stress (ELS) induces long-term vulnerability to psychiatric disorders, particularly those associated with emotional dysregulation, anxiety, and depression (Teicher et al 2016, VanTieghem & Tottenham 2018). The corticolimbic system, including projections from the basolateral amygdala (BLA) and ventral hippocampus (vHIP) to the medial prefrontal cortex (mPFC), plays a pivotal role in the regulation of fear (Giustino & Maren 2015). The prelimbic (PL) region of the mPFC is particularly involved in the process of fear acquisition (Burgos-Robles et al 2009, Corcoran & Quirk 2007, Vidal-Gonzalez et al 2006) and there are strong reciprocal connections between the BLA and mPFC, which associate aversive stimulus to auditory cues during fear conditioning (Little & Carter 2013). The connectivity between the amygdala and the mPFC is also regulated by the vHIP as projections from this region encode contextual representations (Tovote et al 2015). The maturation of the corticolimbic circuitry continues into early adulthood, making it adaptive to environmental cues, but also vulnerable to ELS during critical periods (Drzewiecki & Juraska 2020, Tottenham 2020). For instance, we and others previously demonstrated that ELS produces significant morphological and functional changes in BLA neurons (Guadagno et al 2018b, Malter Cohen et al 2013, Raineki et al 2012), enhancing synaptic plasticity and neuron excitability in pre-adolescent males (Guadagno et al 2020). ELS-exposed animals exhibited altered structural and functional connectivity between the BLA and the mPFC in parallel with elevated fear behavior in adulthood (Guadagno et al 2018a, Honeycutt et al 2020, Junod et al 2019). During maturation of the BLA-mPFC projections, ELS-induced changes in the activity of BLA neurons might exert a “bottom-up” effect on the development and function of the mPFC via glutamatergic projections targeting this region. Indeed, early adversity disrupts dendritic structures, spine density and synaptic plasticity in the PL mPFC of adult males (Baudin et al 2012, Monroy et al 2010, Mychasiuk et al 2012) and increases mPFC innervation from the BLA in pre-adolescent and adult female rats (Honeycutt et al 2020). Currently, the early, pre-puberty mechanisms leading to these adult consequences are still unclear.

Here, we initially tested the hypothesis that ELS would modify the *in vivo* glutamatergic response to fear conditioning in the PL mPFC of adult offspring and that some of these changes would already be detected in pre-adolescence. Long range glutamatergic projections to the mPFC are well segregated between layers in the adult mPFC, where BLA and vHIP afferents target primarily layer II and layer V of the PL mPFC, respectively (Anastasiades & Carter 2021). Such laminar distribution is not entirely mature before adolescence (Bouwmeester et al 2002, Cunningham et al 2002), indicating that ELS might also affect laminar organization within the mPFC. To examine the effects of ELS on the laminar distribution of projections to the PL mPFC, we used retrograde tracing to measure projection density from the BLA, vHIP and mediodorsal thalamus (MDThalamus). In addition, we examined layer-specific ELS-induced presynaptic changes using electrophysiological field recordings of glutamatergic transmission in the PL mPFC.

In addition to changes in the activity and distribution of long-range excitatory inputs, ELS might also disrupt the local inhibitory microcircuits within the mPFC that tightly regulate these projections. Within the mPFC, parvalbumin (PV) and somatostatin (SST) interneurons are predominant (Rudy et al 2011) and they emerge during the neonatal period. Local SST and PV interneurons first appear in the neocortex in the first and second postnatal week, respectively (del Rio et al 1994, Pan et al 2019). ELS has been shown to significantly reduce PV expression in the PL mPFC of adolescent males (PND40) (Grassi-Oliveira et al 2016, Holland et al 2014), suggesting that ELS might alter the developmental trajectory of PV interneurons. SST interneurons also affect the excitation/inhibition balance in the mPFC as they are known to inhibit PV interneurons during conditioned fear acquisition in adults (Courtin et al 2014, Cummings & Clem 2020). Whether ELS alters the participation of specific populations of local inhibitory interneurons on glutamatergic regulation in the pre-adolescent mPFC during fear acquisition is currently unclear. In the current study, we investigated potential ELS-induced changes in mPFC inhibitory tone by performing triple immunostaining of cFos, PV, and SST after fear conditioning in pre-adolescent rats. Our results suggest that ELS modifies both glutamatergic release probability and projection density in a layer specific manner and that activation of PL mPFC PV, but not SST interneurons after fear conditioning is enhanced by ELS in pre-adolescent offspring.

## Methods

### Animals

Timed-pregnant (gestation day 14) Sprague Dawley female rats (Charles River, Kingston, VT) were individually housed under controlled conditions of light (LD 12h:12h, reverse cycle, light on at 8 PM), temperature (21°C-23°C), and humidity (40%-70%), and provided *ad libitum* access to food and water. The day of parturition was considered as postnatal day (PND) 0, and litters were culled to 10 pups on PND1 with both males and females in the litters. Animals were weaned and group housed by sex and treatment on PND21. A total of 63 mothers and their litters were used in these studies distributed as follows: one cohort of 28 dams was used for *in vivo* microdialysis experiments, one cohort of 14 dams was used for immunohistochemistry, one cohort of 12 dams for retrograde tracing experiments, and another cohort of 21 dams was used for electrophysiology experiments. All experimental procedures were reviewed and approved by the Douglas Institute Animal Care Committee at McGill University in accordance with the ethical guidelines from the Canadian Council on Animal Care.

### Limited bedding paradigm

To induce chronic early life stress in the offspring, we used the limited bedding (LB) and nesting protocol adapted from the Baram Lab (Molet et al 2014, Walker et al 2017) and applied between PND1 and 10. On the afternoon of PND1, dams and their offspring were randomly assigned to the normal bedding (NB) or LB condition. In the LB condition, dams and their pups were placed on a metal mesh platform raised 2.5cm above the cage floor and thus elevated from the woodchip bedding on the floor. The mothers were given one-half piece of paper towel as nesting material. The NB cages were given approximately 2 cm layer of woodchips and one piece of paper towel. On PND10, all cages were returned to the NB condition. Mothers and litters were weighed on PND4, PND10, PND14, and PND21 when cages were changed. Maternal behavior (active/passive nursing, pup grooming, self-grooming, eating, drinking, wandering, tail chasing) was video recorded for 24 hours between PND5 and PND6 using infrared cameras and scored using four 60 min observation sessions (two sessions in the dark phase at ZT 14-15, ZT 20-21; and two sessions in the light phase at ZT 2-3, ZT 8-9). Behavior was scored at one-minute intervals during each observation session. A fragmentation of maternal behavior score of 1 was noted when behavior changed from one minute to the next and zero if there was no change.

### In vivo microdialysis

We focused on the right PL mPFC because our previous studies showed that the effects of LB on the synaptic plasticity of BLA neurons and the resting-state BLA-mPFC functional connectivity were more pronounced in the right hemisphere (Guadagno et al 2020, Guadagno et al 2018b). <u>Surgery:</u> A guide cannula used to insert the microdialysis probe into the PL mPFC target site was implanted in pre-adolescent (PND23-25) and adult (PND61-62) rats of both sexes and originating from either NB or LB conditions. Isoflurane (2%-4%) anesthetized animals were stereotaxically implanted with a 22-gauge stainless steel guide cannula (Plastics One, Roanoke, USA) into the right PL mPFC at the following coordinates taken from the Paxinos atlas (Paxinos & Watson 2006): Pre-Adolescent: anteroposterior (A/P) +2.8mm anterior to bregma, lateral (L) +0.51mm right to the midline, and dorsoventral (D/V) 2.3mm below the skull surface; Adults: A/P 3.1mm anterior to bregma, L +0.6mm right to the midline, and D/V 2.5mm below the skull surface. The cannula was secured with acrylic dental cement anchored by two screws threaded into the cranium. A dummy head cap extending 2.7 mm beyond the bottom tip of the cannula (total length: 5.0 mm for pre-adolescents, 5.2 mm for adults) was inserted to prevent infection, cerebrospinal fluid (CSF) seepage, and to accommodate space for the probe on the experimental day. The skin incision was sutured with surgical silk and all rats received 0.5 ml of subcutaneous (s.c.) 0.9% saline for hydration. All animals were allowed a minimum recovery of 5 days before testing. Adult rats were singly housed prior to microdialysis testing and juveniles were pair housed with a perforated Plexiglas barrier dividing the housing space in the cage, allowing smell and sight between the two compartments and thus minimizing social isolation distress.

<u>Microdialysis probes:</u> We used home-made I-shaped microdialysis probes (Luczynski et al 2015) comprised of side-by-side fused quartz inlet and outlet (internal diameter [ID] 50 μm) wrapped in polyethylene tubing (ID 0.58-0.38mm). A regenerated, hollow cellulose membrane (Spectrum, molecular weight cut-off 13 kD; OD 216 μm; ID 200 μm) was secured to the end of a stainless-steel cannula (26 gauge) using cyanoacrylate adhesive and was sealed at its tip with Epoxy. The length of the active membrane was 1.5 mm. On the day of testing, the probe was inserted into the guide cannula, and the probe assembly was attached to a stainless-steel spring that was connected to a liquid swivel (CMA Microdialysis). A computer-controlled microdialysis pump (CMA Microdialysis) was used to pump artificial CSF (aCSF: 26 mM NaHCO_3_, 3 mM NaH_2_PO_4_, 1.3 mM MgCl_2_, 2.3 mM CaCl_2_, 3.0 mM KCl, 126 mM NaCl, 0.2 mM L-ascorbic acid, pH=7.2) through the probe during microdialysis at a rate of 1µl/min. The dialysate was collected at 10 min intervals (10µl) from the quartz outlet. The dead volume of the system was approximately 5 µl.

*<u>In vivo</u>* <u>microdialysis during fear conditioning:</u> *In vivo* microdialysis was performed on PND28-32 in pre-adolescent rats and on PND67-70 in adult rats. Testing took place in fear conditioning chambers (Actimetrics) in semi-dark conditions to respect the reverse light:dark cycle. A red lamp was placed on top of the cage to allow for video recordings of freezing behavior during testing. The fear chamber was cleaned with Peroxigard between trials. Animals were habituated to the fear conditioning chamber without cues or stimuli for 10 min per day on two consecutive days before testing. On the day of testing, a microdialysis probe was inserted into the animals’ pre-implanted guide cannula and the probe was perfused with sterile, degassed aCSF at a constant flow rate of 1µl/min. Dialysate samples were collected but discarded during the first hour of testing to habituate the animals to the microdialysis context and the presence of the experimenter. Baseline microdialysate samples (10µl) were collected every 10 min for 60 min before exposure to a 40 min session of fear conditioning. During fear conditioning, animals were exposed to ten tone (50dB, 30 sec)-shock pairings (0.5sec, shock of 0.5mA, co-terminating with the tone) with an average of 4 min variable inter-trial interval. Recovery dialysate samples were collected for 80 min after the end of fear exposure. Dialysate samples were collected in microtubes containing 1µl of 0.125 M perchloric acid to prevent dialysate degradation, and were immediately frozen at -80°C prior to High Pressure Liquid Chromatography (HPLC) analysis. Freezing behavior was video recorded before, during, and after fear conditioning and freezing behavior during the tones and inter-trial intervals of fear conditioning was manually scored and converted to percentage of freezing time. Freezing was defined as the absence of movement, except for respiration (Stevenson et al 2009). All animals were sacrificed at the end of the microdialysis session and brains were extracted for histological identification of probe placement.

<u>Histology:</u> Microdialysis probe placement was confirmed from 20 µm coronal mPFC brain sections stained with cresyl violet. Only animals with correct placement in the PL mPFC were included in the analysis. The total number of animals included were: pre-adolescent: 30 males and 21 females, adults: 27 males and 22 females.

### Detection of glutamate concentrations by high performance liquid chromatography (HPLC)

Extracellular concentrations of glutamate in the dialysate were quantified by high-performance liquid chromatography with fluorescence detection (HPLC-FD) as previously described (Luczynski et al 2015). The chromatographic system was composed of a pump (UltiMate 3000 RS Pump, Dionex) and an injector connected to an Xterra MS C18 3.0mm x 50 mm, 5 µm analytical column (Waters Corp.). The mobile phase consisted of 3.5% acetonitrile, 15% methanol, and 100 mM Na_2_HPO_4_ and was adjusted to a pH of 6.7. The flow rate was set at 0.5ml/min, and the fluorescence detector (UltiMate 3000 Fluorescence Detector, Dionex) was set to an excitation frequency of 323 nm and to an emission frequency of 455 nm. On the day of the HPLC assay, the dialysate samples were transferred to fraction vials maintained on ice. Working standards (100 ng/ml glutamate) and derivatization reagents were prepared freshly. Standards were loaded with the samples into a refrigerated (10°C) autosampler (UltiMate 3000 RS Autosampler, Dionex). Before being injected into the analytical column, each sample was sequentially mixed with 20 µl of o-phthalaldehyde (OPA, 2.85 mM) diluted with 0.1M Na B O and 20 µl of 3-mercaptopropionic acid (3-MPA, 75mM) diluted with H_2_O and left to react for 5 min. After each injection, the injection loop was flushed with 20% methanol to prevent contamination of subsequent samples. The derivatization reagents (OPA and 3-MPA) were refilled every 48 h until the end of a run. Under these conditions, the retention time for glutamate was approximately 0.7-1 min, with a total run time of 24 min/sample. Chromatographic peak analysis was performed by identification of unknown peaks in a sample according to retention times from known standards (Chromeleon Chromatography Data System software version 7, ThermoFisher Scientific). The glutamate concentrations in the samples collected during and after fear conditioning were normalized to the average of six baseline samples from each individual.

### Electrophysiological field recordings on mPFC slices

<u>Slice preparation for field recording:</u> To evaluate glutamate transmission at the synaptic level, we performed electrophysiological field recordings of the PL mPFC in pre-adolescent NB or LB males aged PND28-35 (n=8-15 animals per group). Animals were anesthetized with a ketamine-xylazine cocktail (0.1 ml/100g body weight, s.c.) and transcardially perfused with ice-cold NMDG-substituted aCSF (in mM: 92 NMDG, 2.5 KCl, 1.25 NaH_2_PO_4_, 30 NaHCO_3_, 20 HEPES, 25 glucose, 2 thio-urea, 5 Na-ascorbate, 3 Na-pyruvate, 0.5 CaCl_2_·2H_2_O, 10 MgSO_4_·7H2O, pH 7.4) until the liver became pale yellow, as described before (Ting et al., 2014). The brain was rapidly extracted, marked on the left hemisphere with a blade, and the anterior brain containing the right mPFC was sliced into 350 µm coronal sections using a Vibratome (Leica Microsytstems) in the same solution. After sectioning, brain slices were incubated in NMDG aCSF maintained at 32°C for 10 min and then kept in HEPES holding aCSF (in mM: 92 NaCl, 2.5 KCl, 1.25 NaH_2_PO_4_, 30 NaHCO_3_, 20 HEPES, 25 glucose, 2 thio-urea, 5 Na-ascorbate, 3 Na-pyruvate, 2 CaCl_2_·2H_2_O, 2 MgSO_4_·7H_2_O, pH=7.4) at room temperature for at least 1 hour before recording. NMDG aCSF and holding HEPES aCSF were freshly prepared from 10x stock and continuously oxygenated with a mixture of 95% O_2_/ 5% CO_2_.

*<u>Ex vivo</u>* <u>electrophysiological field recording:</u> Brain slices including the right mPFC were placed in a recording chamber perfused with oxygenated aCSF (in mM: 125 NaCl, 2.5 KCl, 2 CaCl_2_, 2 MgCl_2_, 1.25 NaH_2_PO_4_, 26 NaHCO_3_, 25 glucose, pH=7.4, Osm 310-320). Evoked field excitatory postsynaptic potentials (fEPSPs) were induced by a bipolar stimulating electrode (#30255, Frameless Hardware Company) and recorded via an aCSF-filled glass electrode. To mainly target BLA afferents to PL mPFC, the stimulating electrode was positioned at the layer I/II border of the right PL mPFC, while the recording electrode was placed in layer II/III of the right PL mPFC. To target vHIP afferents, the stimulating electrode was placed at the border of layer V and VI, with the recording electrode in layer V. The intensity of electrical stimulation for paired-pulse experiments was adjusted to the stimulation intensity that evoked the 40% maximum amplitude of field potential response. Paired pulses were recorded with various intervals at 25ms, 50ms, 100ms, and 200ms, and at least 10 traces were recorded for each interval. For each slice recording, the paired-pulse ratio (PPR) was calculated by dividing the slope of the second fEPSP response by that of the first response, taking the slope value between 10-60% of fEPSP responses. All recordings were amplified by MultiClamp 700B and stored in a PC for offline analysis using Clampfit software (Axon, Molecular Devices).

### Retrograde tracing of right PL mPFC projections

In order to determine the proportion of projections to the PL mPFC from the BLA, vHIP and mediodorsal thalamus (MDThal), pre-adolescent and adult NB and LB male offspring (5-7 rats/group) were injected on PND21 or PND61 with a fluorescent retrograde tracer (Cholera toxin B, CTb, #C34778, ThermoFisher, Canada) in the right PL mPFC (Layer II/III or Layer V). Isoflurane (2%-4%) anesthetized animals were placed in a stereotaxic apparatus and a small hole was drilled in the skull to lower a CTb glass-filled pipet at the following coordinates: Pre-adolescent: Layer II/III: A/P Bregma +2.9 mm, L +0.55 mm, D/V -2.6 mm; Layer V: A/P Bregma +2.9 mm, L +0.95 mm, D/V -2.6 mm; Adult: Layer II/III: A/P Bregma +3.15 mm, L +0.65 mm, D/V -2.95 mm; Layer V: A/P Bregma +3.15 mm, L +1.15 mm, D/V -2.95 mm. CTb (40 nl in pre-adolescents and 80 nl in adults) was pressure-injected in 4 or 8 intervals (10 nl each) through a glass pipet and the pipet was left in place for 5 min before removing it from the brain. Animals were weaned and housed in pairs according to neonatal treatment (NB or LB) and perfused transcardially 1 week after injection with ice-cold 0.9% saline for 5 min, followed by a 20 min perfusion with 4% paraformaldehyde (PFA) in 1X Phosphate Buffer (18.98 mM NaH_2_PO_4_·H_2_O, 95.8 mM Na_2_HPO_4,_ pH=7.41). The brains were extracted and stored in 4% PFA at 4°C overnight, then transferred to a 30% sucrose solution in 1X phosphate-buffered saline (PBS) (137 mM NaCl, 2.7 mM KCl, 10 mM Na_2_HPO_4,_ 1.8 mM KH_2_PO_4,_ pH=7.4) for 48 h at 4°C. The left side of the brains was marked using a blade, and brains were stored at -80°C until slicing. Twenty µm coronal sections of the mPFC were collected on glass slides and imaged under a fluorescent microscope (Zeiss Observer Z1) to assess correct placement of the CTb injection. Fifty µm coronal sections including target areas projecting to the mPFC were collected and stored on uncharged slides at -20°C until being processed for CTb imaging. Images of BLA (anterior and posterior), vHIP and MDThal sections were taken with a Zeiss Observer Z1 fluorescence microscope. Z-stack images of the regions of interest (ROIs) were acquired with a step size of 3µm. The images were taken at 20x magnification, with exposure time of 30ms for DAPI (4’,6-diamidino-2-phenylindole) and 1s for CTb (Far red). The number of CTb-positive cells was counted manually and expressed as density of cells/surface analyzed (mm^2^). Three to five sections per animal were analyzed for each ROI.

### Fear-induced interneuron activation in the PL mPFC

To examine whether fear-induced activity of mPFC PV and SST interneurons was modified by early life experience, we performed triple fluorescence immunohistochemistry for Fos, SST and PV on sections of the PL mPFC from brains of <u>p</u>re-adolescent (PND28-29) offspring exposed to fear conditioning using a protocol similar to the microdialysis experiments. Briefly, 23 pre-adolescent (PND28-29) males from either NB or LB mothers were assigned to either a fear conditioning group or a naïve control group (n=5-6 rats per group). Fear conditioning boxes were the same as those used for microdialysis experiments and animals were tested in semi-dark conditions under red light. In the fear group, animals were placed in the chambers, given 30 min to acclimatize to the experimental context before they were exposed to a 40 min fear conditioning session using the same parameters described above for microdialysis. The “fear” condition can be constructed as a composite of novel environment and shock exposure, similarly to the conditions used in our microdialysis experiments. Animals were perfused 60 min after the onset of fear conditioning. Naïve animals were kept undisturbed in their home cage until anesthetized for perfusion and tissue collection. Animals were perfused and brains were fixed as described above (see retrograde tracing experiments) for determination of Fos, PV and SST by immunohistochemistry.

<u>Triple fluorescence immunohistochemistry for Fos, PV and SST:</u> Fifty µm free floating sections of the mPFC were brought to room temperature for 30 min and washed 3 x 5 min in 1X PBS. They were incubated for 20 min with 0.3% H_2_O_2_ (30%, H1009, Sigma Millipore) in 1X PBS, then washed 3 x 5 min in 1X PBS. After 1 h incubation in blocking solution (3% Normal Goat Serum, S-2000, Vector Laboratories; 0.25% Triton X-100, Sigma Millipore; 1X PBS), sections were incubated with the primary rabbit anti-Fos (1:2000, #226008, Synaptic System) and mouse anti-SST (1:500, sc-74556, Santa Cruz) antibodies for 45 min at room temperature followed by overnight incubation at 4°C on a rotating platform. The following day, sections were washed 3 x 5 min in 1X PBS and incubated for 2 h with the Secondary Goat Anti-Rabbit antibody Alexa-488 (1:1000, A11008, Invitrogen by ThermoFisher Scientific) and Goat Anti-Mouse antibody Alexa-568 (1:1000, A11031, Invitrogen by ThermoFisher Scientific) at room temperature. All procedures were performed in the dark from this point on. The sections were washed 3 x 5 min in 1X PBS and then incubated with the primary Guinea Pig anti-PV antibody (1:2000, #195004, Synaptic Systems) for 45 min at room temperature, then overnight at 4°C. The next day, sections were washed 3 x 5 min in 1X PBS and then incubated with the Secondary Goat Anti-Guinea Pig antibody Alexa-647 (1:1000, #106-605-003, Jackson ImmunoResearch) for 2 h. Sections were washed 3 x 5 min in 1X PBS and mounted onto charged slides using DAPI Hardset mounting medium (H-1500, Vector Laboratories). The slides were stored at 4°C in the dark until imaging.

<u>Microscopy imaging and cell quantification:</u> Images of brain sections containing cFos/PV/SST were taken with a Zeiss Observer Z1 fluorescence microscope. Z-stack images of the PL mPFC were taken with a step size of 3µm. The images were acquired at 20X magnification with an exposure time of 50ms for DAPI, 120ms for GFP (Fos), 220ms for RFP (SST), and 67ms for FR (PV) in triple immunostaining series. The quantification of Fos, SST, and PV-positive cells as well as co-expression of Fos in SST and PV positive cells was performed manually and divided into layer II/III and layer V. The PL mPFC area of cell quantification was outlined with µm as unit of length and total cell counts were converted to cell density measurement in mm^2^. For each animal included, 3-5 brain sections were analyzed.

### Statistical analysis

All data were reported as mean (+/- SEM). Body weight data were analyzed using a two-way ANOVA with bedding as a between-subject factor and postnatal day as a within-subject factor. Two-way ANOVA was also performed on maternal behavior with bedding as between subject factor and light phase as within subject factor. In microdialysis experiments, we first used an unpaired two-tailed Student’s t-test to test for bedding group differences in averaged baseline glutamate levels. Since there were no differences in baseline levels, microdialysis data were expressed as a function of baseline and analyzed using a two-way ANOVA with bedding as a between-subject factor and time as a within-subject factor. Post-hoc Dunnett’s tests were used to compare the experimental timepoints against the baseline. In the microdialysis data, extreme outlier values for the baseline period that were more than +/- 1.5 SD were removed (27 values out of 564). The relatively liberal exclusion approach was applied because the sample size of baseline (n=6) was small in each animal compared to the remainder of the samples (n=12). Outlier values in the fear conditioning and recovery sessions that exceeded 2 SD were excluded from the analysis. Missing and excluded outlier values in the mixed design two-way ANOVA analysis of microdialysis data were estimated using the formula *X*. = [r*U* + *β* (*AB*ij) – *A*i]/[(r − 1)(*β* − 1)] as described previously (Cochran & Cox, 1957). Behavioral freezing responses were analyzed with a two-way ANOVA using bedding as a between factor and time as a within factor. Field recording data (PPR of fEPSP slopes) and the effects of bedding condition and interval were analyzed with two-way ANOVAs with bedding condition as a between-subject factor and interval as a within-subject factor. Density of CTb-positive cells in the various ROIs were analyzed using a two-way ANOVA with bedding and injection sites (layer) as between-subject factors. For the mPFC immunohistochemical data, the effects of LB on the expression of Fos, PV, SST, and colocalization of Fos with either PV or SST were analyzed using a two-way ANOVA with bedding and treatment (naïve vs fear) as between-subject factors. Significant interactions were further assessed by simple main effects analysis, followed by post-hoc Bonferroni’s tests to compare bedding conditions and treatments. For all analyses, the level of significance was set at p<0.05. Graphs were created and statistical analyses were performed with Prism 10 (GraphPad Software).

## Results

### Effect of limited bedding (LB) on maternal behavior and offspring body weight

Exposure to limited bedding between PND1-10 was used to induce early stress in the offspring. Accordingly, pup body weight calculated as the average litter weight was significantly lower in LB compared to NB offspring (Table 1). Two way ANOVA with repeated measures across age showed significant effects of age (F(1.306, 47)=1416, p<0.0001) and bedding (F(1,36)=8.143, p=0.0071) as well as a significant interaction between age and bedding (F(4,144)=3.259, p=0.0136). Post-hoc Bonferroni’s test revealed that LB conditions transiently reduced offspring weight on PND 4 (p=0.0414), PND 10 (p<0.0001), and PND 14 (p=0.0232). By PND 21, LB pup body weight was similar to control NB conditions, showing no significant difference between the NB and LB groups (p=0.543). Changes in offspring body weight were observed in the face of modest changes in maternal behavior (Table 1).

**Table 1.**
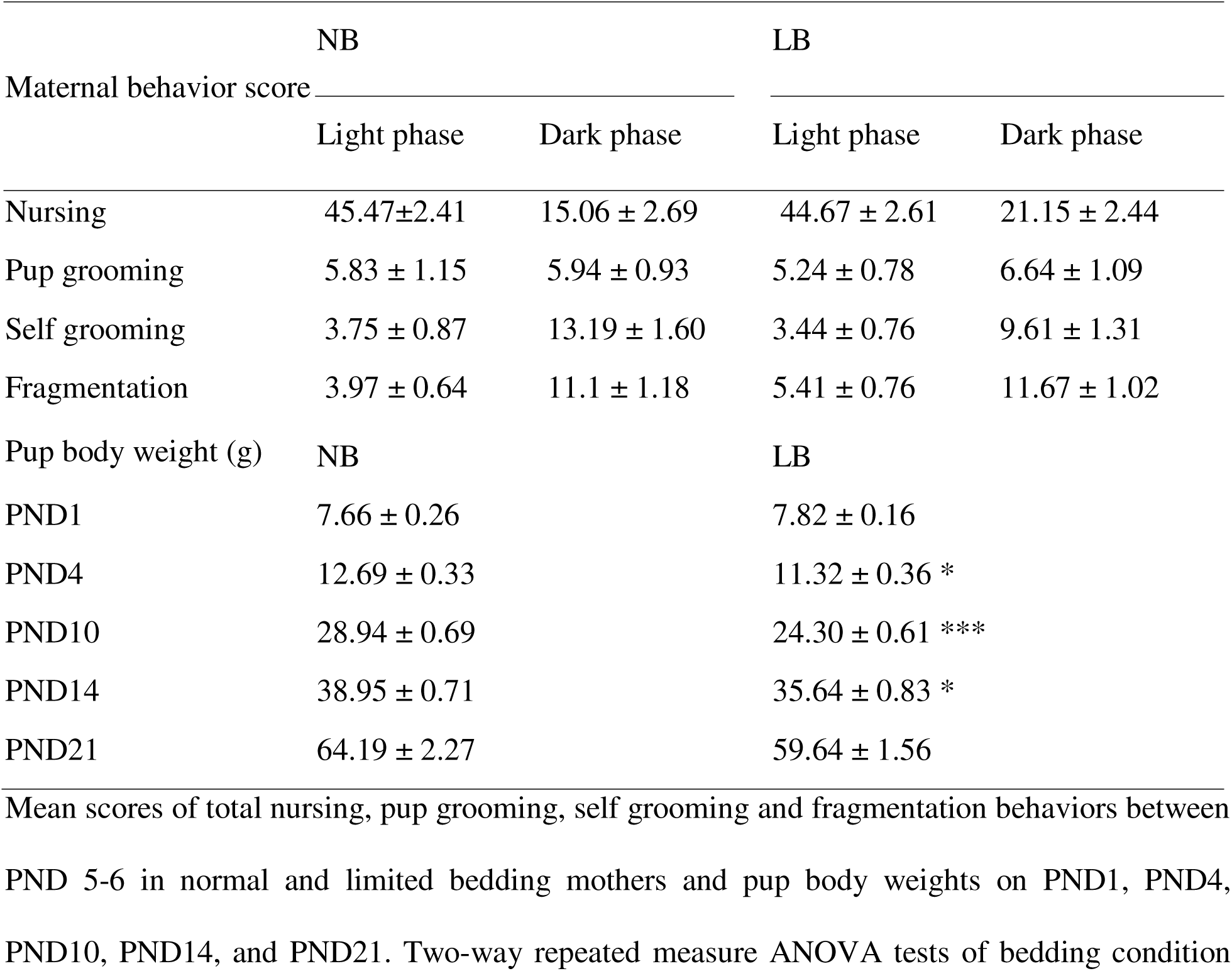
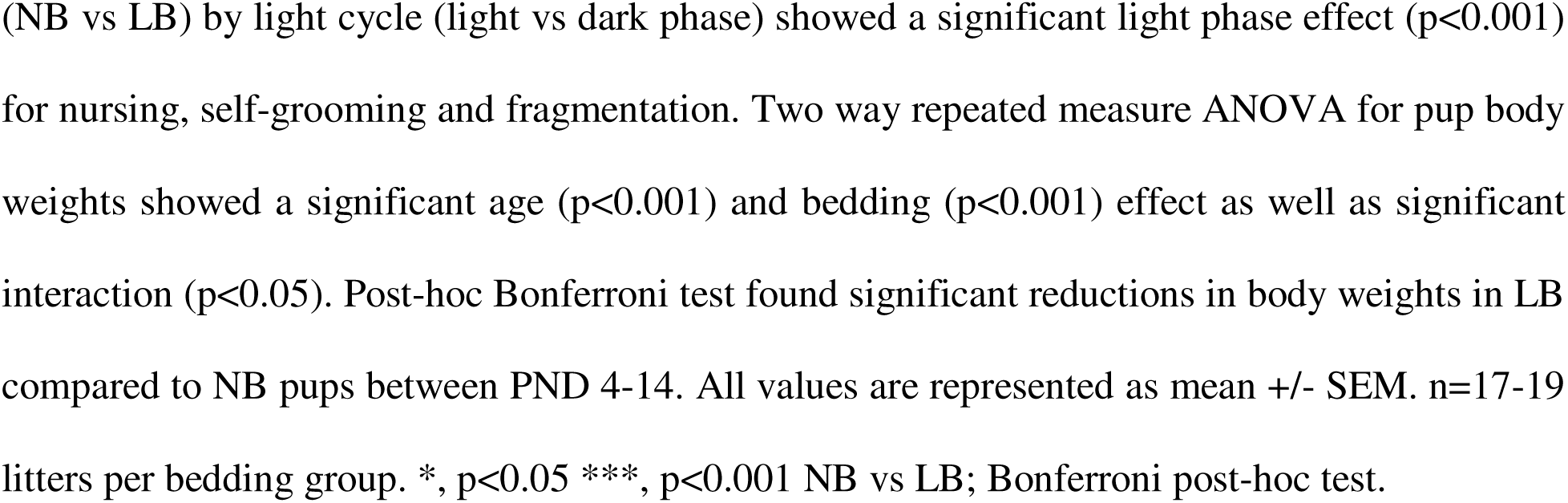
Maternal behavior scores and body weights in normal and limited bedding pups.

| Maternal behavior score | NB |  | LB |  |
| --- | --- | --- | --- | --- |
|  | Light phase | Dark phase | Light phase | Dark phase |
| Nursing | 45.47±2.41 | 15.06 ± 2.69 | 44.67 ± 2.61 | 21.15 ± 2.44 |
| Pup grooming | 5.83 ± 1.15 | 5.94 ± 0.93 | 5.24 ± 0.78 | 6.64 ± 1.09 |
| Self grooming | 3.75 ± 0.87 | 13.19 ± 1.60 | 3.44 ± 0.76 | 9.61 ± 1.31 |
| Fragmentation | 3.97 ± 0.64 | 11.1 ± 1.18 | 5.41 ± 0.76 | 11.67 ± 1.02 |
| Pup body weight (g) | NB |  | LB |  |
| PND1 | 7.66 ± 0.26 |  | 7.82 ± 0.16 |  |
| PND4 | 12.69 ± 0.33 |  | 11.32 ± 0.36 * |  |
| PND10 | 28.94 ± 0.69 |  | 24.30 ± 0.61 *** |  |
| PND14 | 38.95 ± 0.71 |  | 35.64 ± 0.83 * |  |
| PND21 | 64.19 ± 2.27 |  | 59.64 ± 1.56 |  |

Mean scores of total nursing, pup grooming, self grooming and fragmentation behaviors between PND 5-6 in normal and limited bedding mothers and pup body weights on PND1, PND4, PND10, PND14, and PND21. Two-way repeated measure ANOVA tests of bedding condition
(NB vs LB) by light cycle (light vs dark phase) showed a significant light phase effect ( $p < 0.001$ ) for nursing, self-grooming and fragmentation. Two way repeated measure ANOVA for pup body weights showed a significant age ( $p < 0.001$ ) and bedding ( $p < 0.001$ ) effect as well as significant interaction ( $p < 0.05$ ). Post-hoc Bonferroni test found significant reductions in body weights in LB compared to NB pups between PND 4-14. All values are represented as mean ± SEM. $n = 17-19$ litters per bedding group. \*, $p < 0.05$ \*\*\*, $p < 0.001$ NB vs LB; Bonferroni post-hoc test.

A two-way ANOVA, with bedding as between-subject factor and light cycle as within-subject factor, showed no significant interaction between bedding conditions and light phase, nor significant effects of bedding conditions on the four categories of behavior analyzed (nursing, self-grooming, pup grooming and fragmentation). Consistent with previous results (Guadagno et al 2018b), both NB and LB mothers nursed more during the light phase than the dark phase (Light Effect: F(1,33)=99.68, p<0.0001), but displayed more self-grooming behavior (Light Effect: F(1,33)=55.57, p<0.0001) and fragmentation (Light Effect: F(1,33)=69.25, p<0.0001) during the dark phase. Although we did not observe significantly increased fragmentation of maternal behavior in LB conditions, the reduction in body weight in LB offspring was consistent with previous studies(Arp et al 2016, Brunson et al 2005, Guadagno et al 2018b), indicating that pups experienced chronic stress in LB conditions.

### Fear-induced glutamate responses in the prelimbic mPFC and freezing behavior

<u>Adults:</u> To examine the effects of LB on fear-induced glutamate response in the PL mPFC when the fear circuitry is fully developed, we first measured extracellular glutamate concentrations before, during and after a 40 min session of fear conditioning in adult males and females using *in vivo* microdialysis (Figure 1A). As illustrated in Figure 1B, only animals with confirmed probe placement in the PL mPFC were included in the analysis (males: n=13-14, females: n=9-13). Basal levels of glutamate release (average of B1-B6) were similar between NB and LB adult offspring in both males (t(26)=1.037, p=0.3091) (Figure 1C) and females (t(22)=0.2090, p=0.8364) (Figure 1D). All values of glutamate concentrations after the onset of fear conditioning were expressed as a percentage of the individual baseline values. Two-way ANOVAs with bedding conditions as between-subject factor and time as within-subject factor were performed to analyze PL mPFC glutamate release in both sexes. In adult males, LB exposure enhanced the fear-induced glutamate responses with a significant main effect of bedding (F(1,25)=4.884, p=0.0365) and time x bedding interaction (F(12,300)=2.513, p=0.0037), but no significant time effect (Figure 1E). Subsequent post-hoc Dunnett’s tests found that glutamate concentrations significantly increased above baseline during F2 (p<0.05) and after fear conditioning (R6 and R8, p<0.05) in the LB exposed adult offspring exclusively, but not in NB controls. When analyzing glutamate concentrations as a function of the layer where the probe was mainly located, we observed a significant bedding effect for layer V (F(1,12)=7.177; p =0.02), but not when the probe was located in layer II/III or at the border of both layers. In contrast, glutamate concentrations in the PL mPFC during and after fear conditioning were not significantly altered between NB and LB adult females (Bedding: F (1, 20) = 2.161, p=0.1571; Time: F(12, 240)=1.483, p=0.1308; Bedding x Time Interaction: F(12,240)=0.7689, p=0.6822) (Figure 1F).

**Figure 1:**
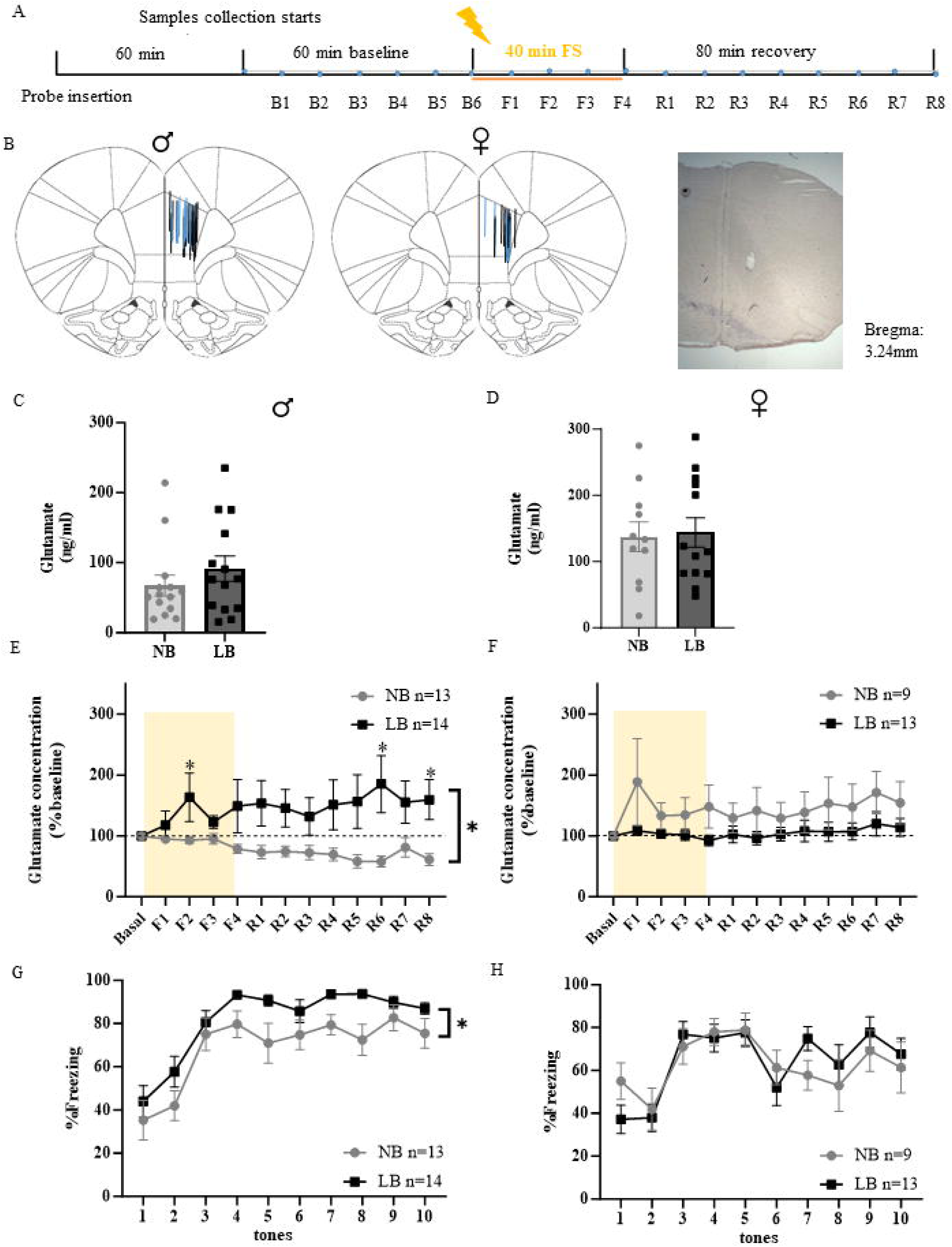
Sex-dependent effects of LB on fear-induced glutamate release in the right PL mPFC of NB and LB adult rats. **A**: Representation of the experimental microdialysis protocol with 60 min of wash time, 60 min of baseline, 40 min of fear conditioning, and 80 min of recovery. Microdialysate samples were collected every 10 min from the start of baseline (B1). **B**: Distribution of the probe placements within the right PL mPFC on the Paxinos atlas (Paxinos and Watson, 2005) and representative coronal brain section for microdialysis probe placement stained with cresyl violet at 4x magnification. Lengths of the blue and black lines correspond to the 1.5 mm length of the active membrane of the microdialysis probes in NB and LB animals, respectively. **C-D**: Basal levels of extracellular glutamate concentrations pooled from B1-B6 samples collected from NB and. LB adult males and females before fear conditioning. There were no significant differences between NB and LB adult offspring in either sex (Student t-test). **E-F:** Extracellular glutamate concentrations normalized as percentage of baseline in the right PL mPFC of NB and. LB adult males and females during (F1-F4) and after (R1-R8) fear conditioning. Shaded areas represent the 40 min duration of fear conditioning. Two-way ANOVA with bedding as between-subject factor and time as within-subject factor showed that normalized fear-induced glutamate concentrations were higher in the PL mPFC of LB adult males compared to NB adult males (bedding effect p<0.05, bedding x time interaction p<0.01), but not in adult females. Glutamate concentrations were significantly increased above baseline during (F2) and after (R6, R8) fear conditioning in the LB adult males (Dunnett’s post-hoc test, p<0.05). **G-H**: Percentage of freezing time during exposure to 10 tone-shock pairings in NB and LB offspring during *in vivo* microdialysis. Although fear conditioning increased freezing behavior in both adult males and females (time effect: p<0.001), bedding effect was significant only in male offspring (p<0.05), where freezing in LB was higher than in NB rats (Two-way ANOVA with bedding as between-subject factor and time as within-subject factor). All values are represented as mean +/- SEM. n=9-14 per group. Figure 1-1: Percentage of freezing time across ten tone-shock intervals during *in vivo* microdialysis in adult male, adult female, pre-adolescent male, and pre-adolescent female offspring. Two-way ANOVAs were conducted with bedding condition as a between-subject factor and interval as a within-subject factor. Fear conditioning significantly increased freezing behavior over ten intervals in all groups of animals (interval effects: p<0.001). A significant main effect of bedding was only observed in adult males (F(1,25)=6.377, p=0.018), where LB animals displayed significantly more freezing compared to NB controls. No significant interaction between bedding x interval was observed. All values are represented as mean +/- SEM. n=9-17 animals per group. *, p<0.05.

Consistent with higher glutamate concentrations in LB adult males during fear conditioning, we also observed increased freezing behavior in LB compared to NB offspring during the 40 min session of fear conditioning (10 tone/shock pairings, Figure 1G). Two-way ANOVAs with bedding conditions as between-subject factor and time as a within-subject factor showed a significant effect of bedding (F(1,25)=7.076, p=0.0134) and time (F(9,225)=21.96, p<0.0001), but no significant interaction in males. In adult females, freezing behavior significantly increased during fear conditioning (Time: F(9,180)=7.716, p<0.0001), but there was no significant difference between NB and LB animals (Bedding: F(1,20)=0.03029, p=0.8636; and no time x bedding interaction(Figure 1H). Freezing behavior during the intervals between the tone shock pairings did not exhibit any effect of bedding for any of the experimental groups tested, except for the adult male group (Figure 1-1), indicating that the behavioral freezing response to context was also higher in LB compared to NB offspring.

<u>Pre-adolescents:</u> Because of the protracted development of the mPFC until early adulthood, we hypothesized that the consequences of early LB conditions on fear-induced glutamate responses might be different between adult and pre-adolescent male and female offspring. In a similar experimental design as for the adults (Figure 2A), we tested rat offspring between PND28-32. This age range was chosen to correspond to a large increase in mPFC glutamatergic projections originating from the BLA and vHIP observed around PND30 (Pattwell et al., 2012). Only animals with PL mPFC probe placement were included in our analyses (Figure 2B, males: n=13-17; females: n=9-12). Similar to the adult offspring, basal levels of glutamate concentrations were not altered by bedding conditions in pre-adolescent males (t(26)=0.6912, p=0.4956, Figure 2C) or females (t(21)=1.239, p=0.2290, Figure 2D). In male offspring, two way ANOVA analysis of PL mPFC glutamate concentrations showed a trend for a significant bedding effect (F(1,28)=2.45, p=0.1287), but no significant time effect (F(12,336)=1.207, p=0.2766), or time x bedding interaction (F(12,336)=1.262, p=0.2395). Even though the bedding effect did not reach significance, glutamate concentrations in the PL mPFC increased after fear conditioning in NB male rats, while this response was absent in LB male rats. Post-hoc Dunnett’s tests revealed that fear conditioning significantly increased PL extracellular glutamate concentrations during the first 20 min of fear conditioning (F1, F2) and the last 10 min of testing (R8) in the NB male offspring only (p<0.05, Figure 2E). This contrasts with the situation we observed in adult offspring, where LB response to fear conditioning was higher than that of NB offspring. On the other hand, similar to adult females, glutamate release in the PL mPFC of pre-adolescent females was not affected by bedding conditions or fear conditioning (Bedding: F(1,19)=0.3773, p=0.5463; Time: F(12, 221)=0.6677, p=0.7815, no interaction) (Figure 2F). Two-way ANOVAs of freezing behavior revealed that both NB and LB pre-adolescent males and females were able to acquire the fear association at a young age (Figure 2G and H), as indicated by increased freezing behavior (Time effect: male F(9,252)=11.09, p<0.0001; female (F(9,171)=9.196, p<0.0001), but the freezing response was not significantly different between bedding conditions and there was no significant time x bedding interaction for either sex. Freezing behavior during the intervals between the tone shock pairings was not significantly affected by bedding (Figure 1-1).

**Figure 2:**
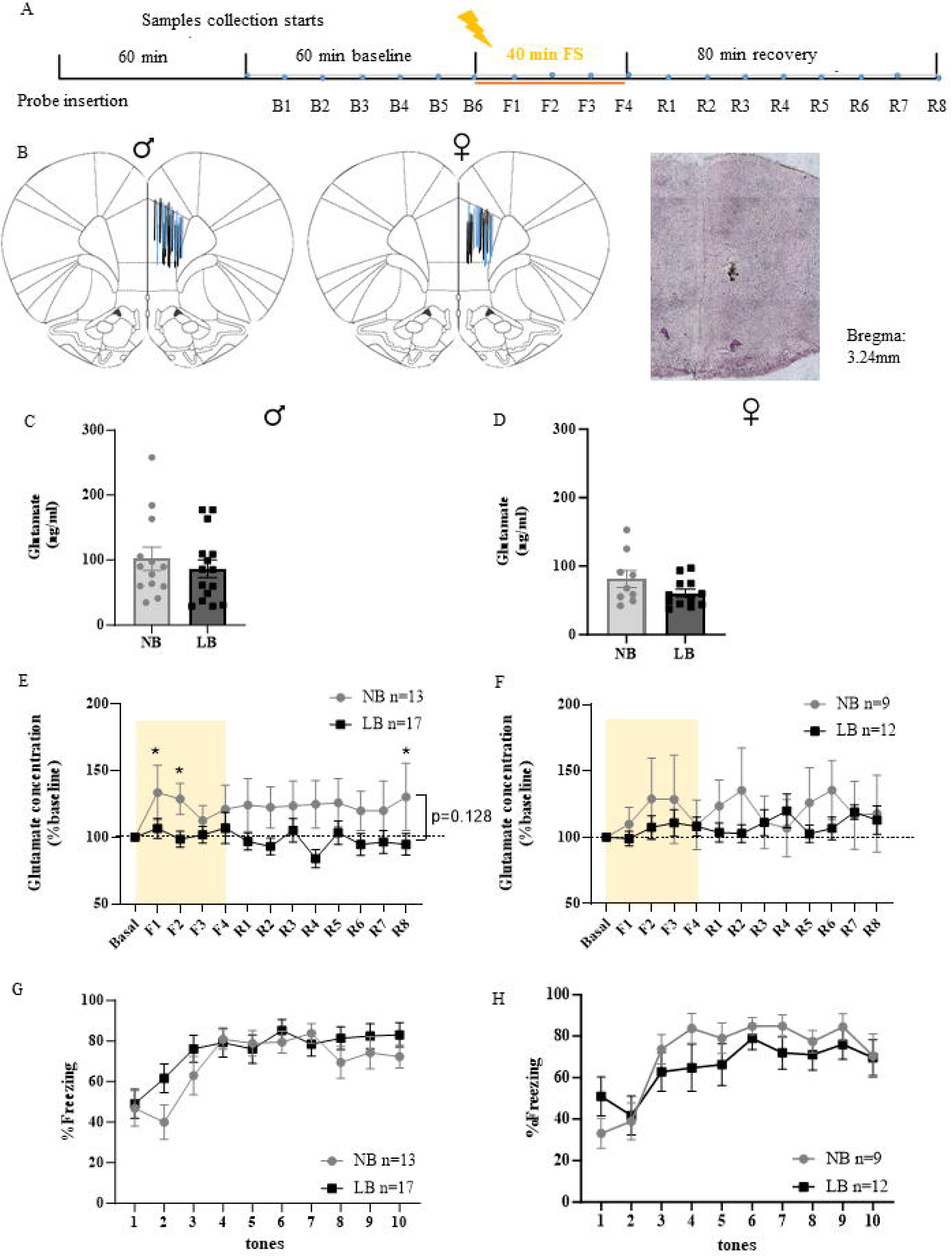
Sex-dependent effects of LB on fear-induced glutamate release in the right PL mPFC of NB and LB pre-adolescent rats**. A**: Representation of the experimental microdialysis protocol with 60 min of wash time, 60 min of baseline, 40 min of fear conditioning, and 80 min of recovery. Microdialysate samples were collected every 10 min from the start of baseline (B1). **B**: Distribution of the probe placements within the right PL mPFC on the Paxinos atlas (Paxinos and Watson, 2005) and representative coronal brain section for microdialysis probe placement stained with cresyl violet at 4x magnification. Lengths of the blue and black lines correspond to the 1.5 mm length of the active membrane of the microdialysis probes in NB and LB animals, respectively. **C-D**: Basal levels of extracellular glutamate concentrations pooled from B1-B6 samples collected from NB and. LB adult males and females before fear conditioning. There were no significant differences between NB and LB pre-adolescent offspring in either sex (Student t-test). **E-F:** Extracellular glutamate concentrations normalized as percentage of baseline in the right PL mPFC of NB and LB pre-adolescent males and females during (F1-F4) and after (R1-R8) fear conditioning. Shaded areas represent the 40 min duration of fear conditioning. Two-way ANOVA with bedding as between-subject factor and time as within-subject factor showed that normalized fear-induced glutamate concentrations tended to be lower in the PL mPFC of LB-exposed pre-adolescent males (p=0.128), but this trend was not observed in females. In males, significant increases in glutamate concentrations over baseline were significant at F1, F2, and R8 in NB offspring (p<0.05). **G-H**: Percentage of freezing time during exposure to 10 tone-shock pairings in NB and LB offspring during *in vivo* microdialysis. Fear conditioning increased freezing behavior in both adult males and females (time effect: p<0.001), with no significant effect of bedding conditions (Two-way ANOVA with bedding as between-subject factor and time as within-subject factor). All values are represented as mean +/- SEM. n=9-17 per group. Figure 1-1: Percentage of freezing time across ten tone-shock intervals during *in vivo* microdialysis in adult male, adult female, pre-adolescent male, and pre-adolescent female offspring. Two-way ANOVAs were conducted with bedding condition as a between-subject factor and interval as a within-subject factor. Fear conditioning significantly increased freezing behavior over ten intervals in all groups of animals (interval effects: p<0.001). A significant main effect of bedding was only observed in adult males (F(1,25)=6.377, p=0.018), where LB animals displayed significantly more freezing compared to NB controls. No significant interaction between bedding x interval was observed. All values are represented as mean +/- SEM. n=9-17 animals per group. *, p<0.05.

In summary, the effects of LB on fear-induced PL mPFC glutamate concentrations were sex-and age-dependent. Fear-induced glutamatergic transmission was elevated by LB exposure in adult males, while it tended to be reduced in this region in pre-adolescent males. No significant effects were observed in females at either age. Because significant LB effects were found mostly in males, our subsequent studies focused on the investigation of male PL mPFC function and potential mechanisms leading to the observed changes. However, we are aware that the lack of LB effect in females might correspond to different developmental processes affected in females that could compensate or provide greater adaptability to early adverse conditions in infancy.

### Effects of LB on layer-specific presynaptic glutamate release in the PL mPFC of pre-adolescent male offspring

In order to investigate whether the reduced glutamate release during fear conditioning in LB-exposed pre-adolescent males might be due to low presynaptic glutamate release in either layer II/III or layer V, we examined *in vitro* presynaptic glutamate transmission by electrophysiological fEPSP recordings on mPFC slices of preadolescent male NB and LB offspring on PND28-35. We recorded fEPSPs that were evoked by paired-pulse stimulations at different inter-pulse intervals (25-200 ms) in either layer II/II or layer V. As illustrated in Figure 3A, and 3D, the stimulating and recording electrodes were placed either in layer II/III or layer V of the PL mPFC. The PPRs expressed as the calculated ratio of the second pulse over the first pulse of fEPSP slopes were analyzed to evaluate presynaptic glutamate release probability (Layer II/III: Figure 3B-C; Layer V: Figure 3E-F). For Layer II/III of the PL mPFC, a two-way ANOVA test with bedding condition as a between-subject factor and interval as a within-subject factor revealed that PPRs were significantly reduced by the LB condition (bedding: F(1,26)=6.561, p=0.0166), but not affected by the interval (F(3,78)=1.434, p=0.2392). There was no bedding x interval interaction for recordings in layer II/III (Figure 3C). When field recordings were conducted with stimulating and recording electrodes placed in the layer V of the PL mPFC, a different pattern emerged. A two-way ANOVA test with bedding condition as a between-subject factor and interval as a within-subject factor revealed that PPRs in the layer V of the PL mPFC were significantly higher in the LB offspring compared to the NB controls (bedding: F(1,18)=6.897, p=0.0171; interval: F(3,54)=2.268, p=0.0910; bedding x interval: F(3,54)=1.766, p=0.1645, Figure 3E-F), suggesting that LB exposure reduced presynaptic glutamate release probability in the layer V of PL mPFC. Collectively, we found that the LB-induced alterations in glutamatergic transmission in the PL mPFC of pre-adolescent males were layer specific, where the probability of presynaptic glutamate release was increased in layer II/III, but decreased in layer V of PL mPFC.

**Figure 3:**
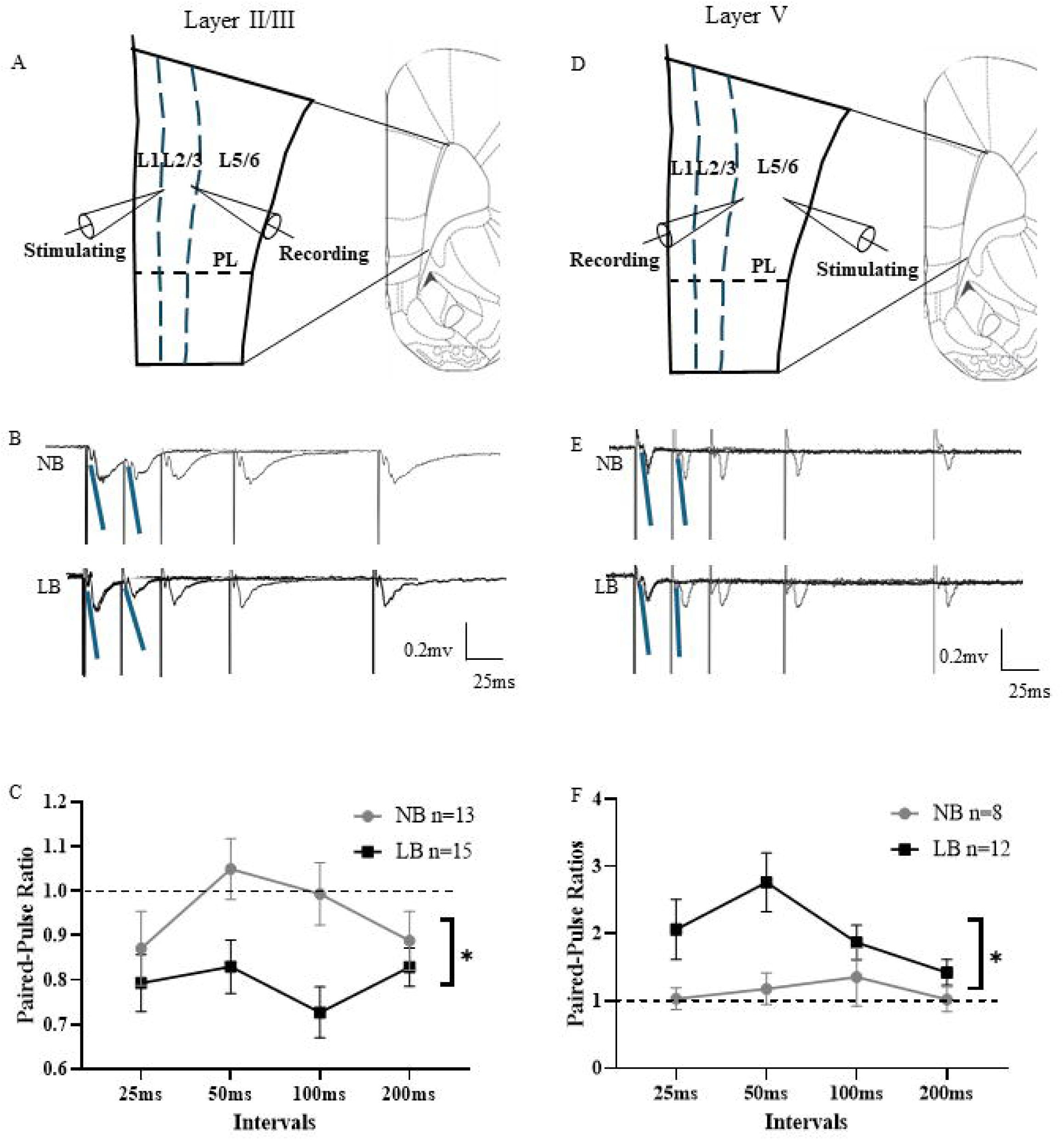
*In vitro* layer specific presynaptic glutamate transmission in slices from the right PL mPFC of NB and LB pre-adolescent male offspring. **A:** Schematic placement of the stimulating and recording electrodes in the layer II/III of PL mPFC. **B**: Representative fEPSP traces evoked by paired pulse stimulations at various inter-stimulus intervals (25-200 ms) in the layer II/III of PL mPFC in NB and LB animals. Blue lines represent the slopes of the paired fEPSPs at 25 ms. (Scale bar: 0.2mV; 25ms). **C**: Paired-pulse ratios (PPRs) of fEPSP slopes at different intervals in the layer II/III of PL mPFC for NB (n=13) and LB (n=15) offspring. Two-way ANOVA with bedding (between-subject) and interval (within-subject) factors showed that PPRs in the layer II/III of PL mPFC were significantly lower in the LB exposed pre-adolescent male offspring compared to NB controls (bedding effect: p<0.05). **D:** Schematic placement of the stimulating and recording electrodes in the layer V of PL mPFC. **E**: Representative fEPSP traces evoked by paired pulse stimulations at various inter-stimulus intervals (25-200 ms) in the layer V of PL mPFC in NB and LB animals. Blue lines represent the slopes of the paired fEPSPs at 25 ms. (Scale bar: 0.2mV; 25ms). **F**: PPRs of fEPSP slopes at different intervals in layer V of the PL mPFC for NB (n=8) and LB (n=12) offspring. Two-way ANOVA with bedding (between-subject) and interval (within-subject) factors showed that PPRs in the layer V of PL mPFC were significantly higher in LB animals than in NB controls (bedding effect: p<0.05). All values are represented as mean +/- SEM. *, p<0.05.

### Layer specific long-range projections to the PL mPFC are altered by LB conditions in pre-adolescent male offspring

To investigate whether layer-specific alterations in PL mPFC presynaptic glutamate release induced by LB might be associated with an altered laminar distribution of long-range glutamatergic projections in this region, we examined the density of projecting neurons in the BLA, vHIP, and MDThal synapsing in either superficial (layer II/III) or deep layers (layer V) of the PL mPFC in NB and LB pre-adolescent males. Besides the BLA and the vHIP, the MDThal is one of the most prominent subcortical regions contributing to glutamatergic afferents to the PL mPFC and participates in fear regulation (Arhlund-Richter et al., 2019; Venkataraman & Dias, 2023). As illustrated in Figure 4A, a retrograde tracer (CTb) was injected in either the layer II/III or the layer V of the PL mPFC on PND21 (left panels) and quantification of CTb+ neurons in the BLA, vHIP, and MDThal was performed a week later (PND28). The anterior and posterior BLA (aBLA: Bregma -1.68 mm to -2.5mm; pBLA: Bregma -2.51mm to 3.36mm) (Paxinos & Watson, 2005) were analyzed separately since LB induces significant differences in projections from these two regions to the mPFC (Guadagno et al., 2018). In the aBLA, a two-way ANOVA test using bedding condition and layer as between-subject factors showed a significant main effect of bedding condition (F(1,19)=5.78; p =0.0266) and a significant interaction between bedding condition and layer (F (1, 19) = 10.59; p =0.0042), without a significant difference between layers (F(1,19)=3.193; p =0.899) (Figure 4B). Bonferroni post-hoc test revealed that LB exposure significantly increased the density of CTb+ neurons in the aBLA that are projecting to layer V (p<0.001), but not layer II/III of the PL mPFC. In LB-exposed offspring, the density of aBLA neurons projecting to layer V was also significantly higher than that projecting to layer II/III of the PL mPFC (p<0.01). The LB-induced changes in PL mPFC projecting neurons in the aBLA of pre-adolescent males did not persist into adulthood (Figure 4-1). Similarly to LB pre-adolescent offspring, in the adult, the density of CTb-positive neurons projecting to layer V of the PL mPFC was significantly higher than that targeting layers II/III (layer effect: F (1, 24) = 12.37, p =0.0018), but there was no significant effect of bedding condition in the adult (Figure 4-1A). These results suggested that the laminar distribution of PL mPFC projecting aBLA neurons was precocious in LB-exposed pre-adolescent male offspring compared to their NB counterparts. In contrast, the density of pBLA neurons projecting to the PL mPFC did not significantly differ between bedding conditions or laminar destination in both young (Bedding: F(1, 20) = 4.290, p =0.0515; Layer: F(1, 20) = 0.6975, p =0.4135; Bedding x Layer Interaction: F(1, 20) = 1.183, p =0.2896; Figure 4C) and adult males (Figure 4-1B).

**Figure 4:**
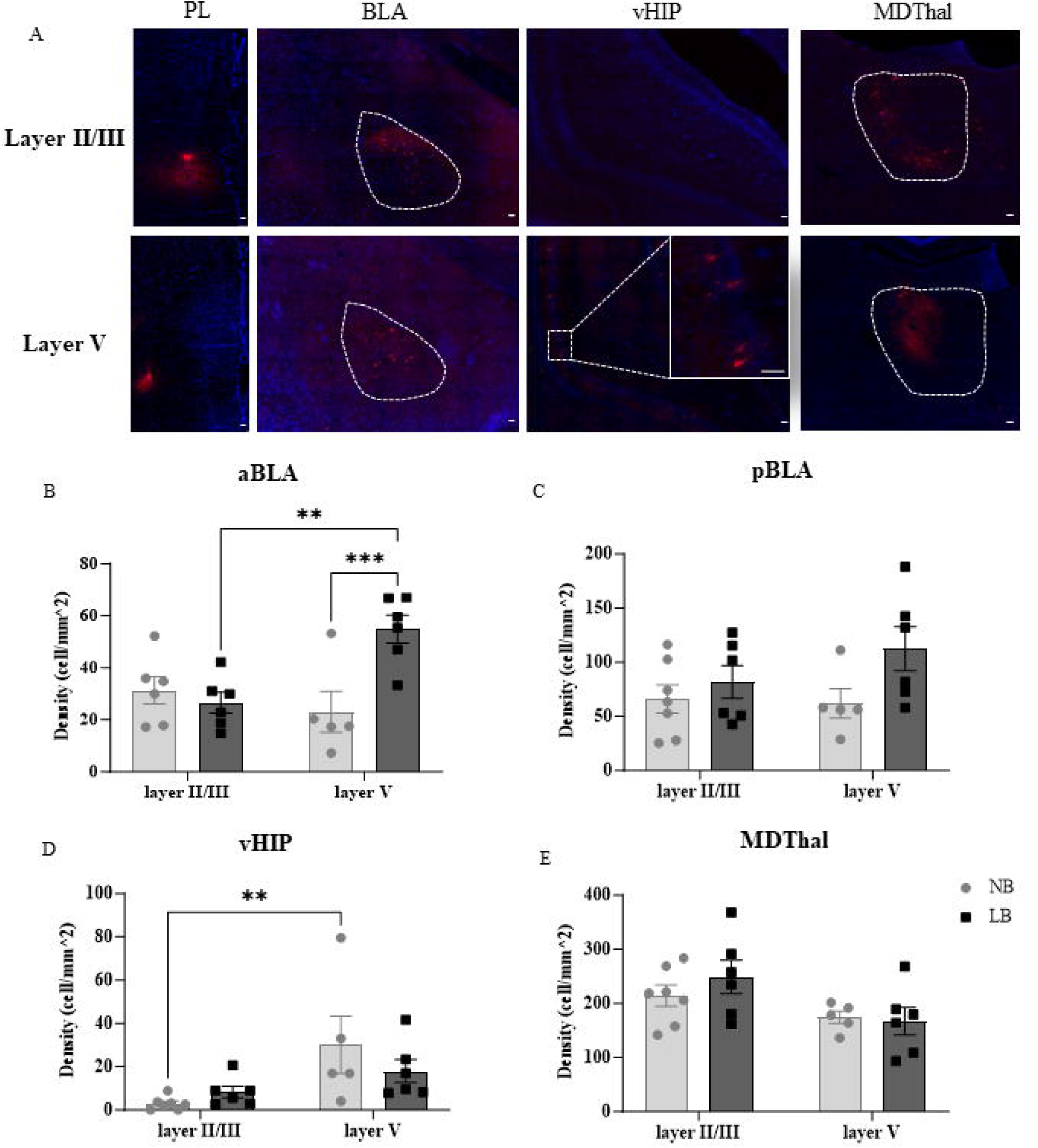
Density of PL mPFC projecting neurons in the right BLA, vHIP, and MDThal of NB and LB pre-adolescent males. **A**: Representative CTb+ fluorescent images taken from the BLA, vHIP, and MD after CTb injection in the layer II/III or layer V of PL mPFC in PND28-29 male NB offspring. Images were taken at 20x magnification and the scale bars represent 50 μm. **B-E**: Density of CTb+ neurons (cells/mm^2^) targeting either layer II/III or layer V of the PL mPFC in the aBLA (B), pBLA (C), vHIP (D) or MD (E) of NB or LB offspring. Two-way ANOVA with bedding and layer as between-subject factors was used to analyze these data. B. Density of CTb+ neurons targeting layer V of the PL mPFC in the aBLA of LB-exposed animals was significantly higher than NB controls (bedding effect: P<0.05; bedding x layer interaction: p<0.01). Compared to layer II/III projecting neurons, there were more aBLA neurons synapsing in layer V in LB-exposed offspring, but not in NB controls. C. Density of PL projecting neurons in the pBLA was not affected by bedding conditions or laminar destinations. D. Density of PL layer V projecting neurons was significantly higher than that of PL layer II/III projecting neurons in the vHIP (layer effect: p<0.01). E. Density of PL Layer V projecting neurons in the MD was significantly lower than that of PL layer II/III projecting neurons (layer effect: p<0.05). All values are represented as mean +/- SEM. n=5-7 animals per group. **, p<0.01. ***, p<0.001. Figure 4-1: Density of PL mPFC projecting neurons in the right BLA of NB and LB adult males. Two way ANOVA with bedding and layer as between subject factors was used to analyze these data. A: Density of CTb+ neurons (cells/mm^2^) projecting to PL mPFC layer V was significantly higher than that targeting layer II/III in the aBLA (layer effect, p<0.01), with no effect of bedding condition. B: Density of PL projecting neurons in the pBLA was not affected by bedding condition or laminar destination. All values are represented as mean +/- SEM, n=4-11 animals per group. **, p<0.01.

In the vHIP, a two-way ANOVA test using bedding condition and layer as between-subject factors showed a significant effect of layer (F(1, 20)=8.733, p=0.0078), but no effect of bedding (F(1,12)=0.2936, p =0.5939) or bedding x layer interaction (Figure 4D). As reported in adults (Liu & Carter, 2018), vHIP projections to the PL mPFC in pre-adolescent rats were higher in layer V compared to layer II/III of PL mPFC. Interestingly, the difference between laminar destinations of vHIP projections in the PL mPFC tended to be significant only in NB (Student t-test, p<0.01), but not in LB exposed animals. In the MDThal, a two-way ANOVA revealed a significant main effect of layer (F(1,20)=6.475, p =0.0193), indicating a higher density of CTb+ neurons when CTb was injected into layer II/III compared to layer V, consistent with previous findings in adults (Anastasiades & Carter 2021). No significant effect of bedding or bedding x layer interaction were observed (bedding: F(1,20)=0.3374, p =0.5678; bedding x layer interaction: F(1,20)=0.7798, p =0.3877; Figure 4E).

### Effect of LB on activation of PL mPFC interneurons after fear conditioning in pre-adolescent male offspring

In the mPFC, glutamatergic inputs not only synapse on excitatory pyramidal neurons, but they are highly gated by the activity of local inhibitory interneurons (Tembley et al. 2016). To test whether reduced fear-induced glutamate concentrations observed in pre-adolescent LB male offspring in our microdialysis experiments could be due to increased inhibitory tone in the PL mPFC, we examined activation of PV and SST interneurons 60 min after the onset of fear conditioning using triple immunostaining for Fos, SST, and PV in the PL mPFC of male offspring on PND28-29 (Figure 5A). We first quantified the density of PV and SST interneurons in the layer II/III and layer V of the right PL mPFC in both NB and LB pre-adolescent males. Two-way ANOVAs were conducted with bedding condition as the between-subject factor and cortical layer as the within-subject factor. The analyses revealed no significant effects of bedding condition or layer on the density of PV+ (bedding effect: F (1, 21) = 0.0007, p =0.9796; layer effect: F (1, 21) = 3.572, p =0.0727; Figure 5B) or SST+ cells (bedding effect: F(1, 21)=1.37, p =0.2556; layer effect: F(1, 21)=0.60, p =0.4464; Figure 5C). Two-way ANOVAs with bedding condition and fear treatment as between-subject factors were used to analyze cFos expression (Figure 5D-E) and co-expression with PV (Figure 5F-G) and SST (Figure 5H-I) in layer II/III and layer V PL mPFC of naïve control and fear-exposed NB or LB offspring. The overall expression of cFos in the PL mPFC was significantly elevated by fear exposure (treatment effect: layer II/III: F(1,19)=18.29 p =0.0004; layer V: F (1, 19) = 20.37, p =0.0002), without a significant bedding effect (layer II/III: F(1,19)=1.366, p =0.257; layer V: F (1, 19) = 0.002695, p =0.9591) or bedding x treatment interaction (layer II/III: F(1,19)=0.7328 p =0.405; layer V: F (1, 19) = 1.176, p =0.2918) in both layers of PL mPFC (Figure 5D-E).

**Figure 5:**
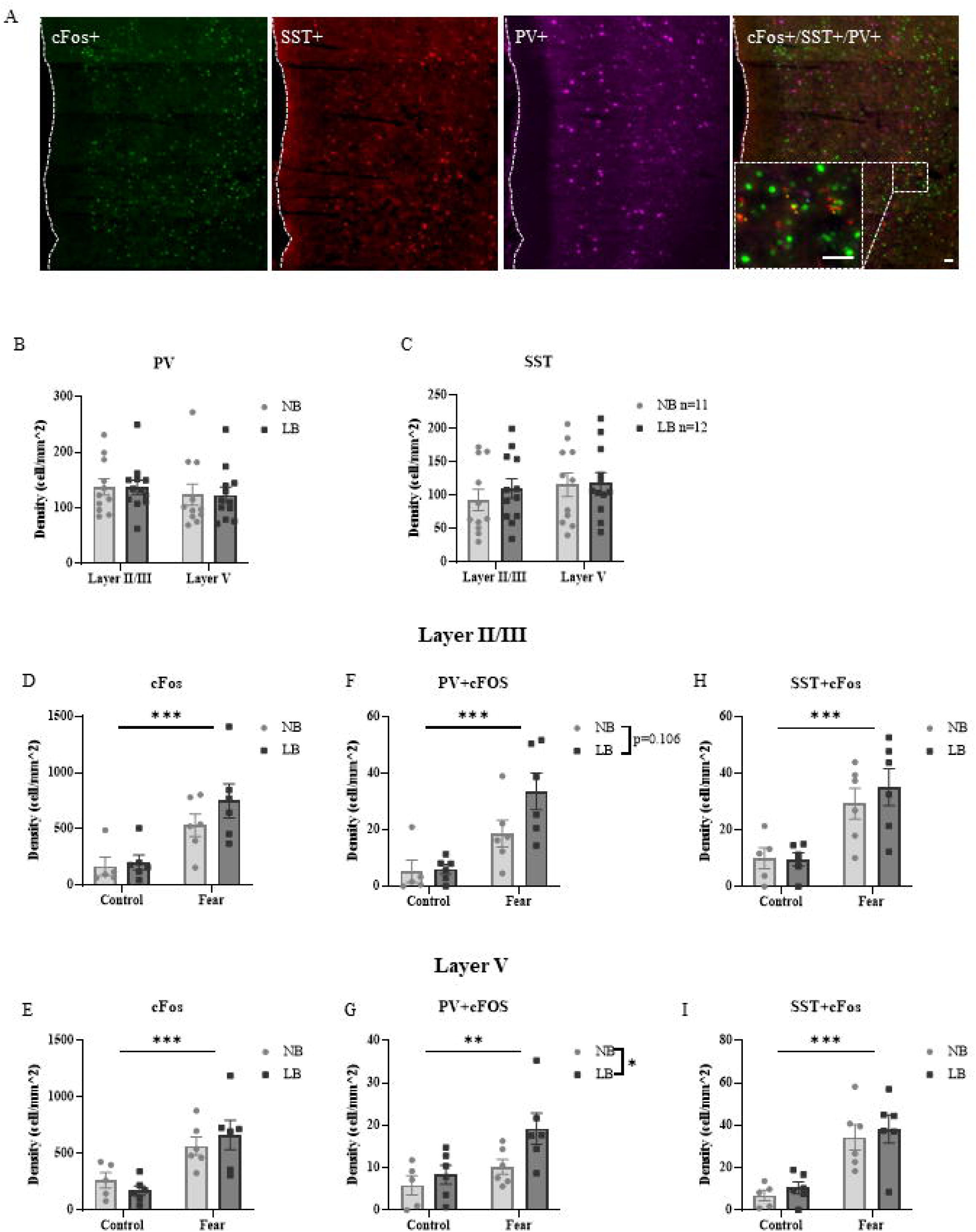
Density and activation of parvalbumin (PV) and somatostatin (SST) interneurons after fear conditioning in pre-adolescent NB and LB males. **A**: Representative triple immunofluorescence staining images taken from the right PL mPFC after fear conditioning in PND28-29 NB male rats showing cFos positive (green), SST positive (red), PV positive (magenta), and merged expression of cFos, SST and PV positive cells. Images were taken at 20x magnification, white dotted line represent the midline and scale bars represent 50 μm. Yellow stars and blue stars indicate the co-localization of SST/cFos and PV/cFos in the zoomed-in image, respectively. Two-way ANOVA with bedding as between-subject factors and layer as within-subject factors were used to analyze the density of PV+ (**B**) and SST+ (**C**) cells in the right PL mPFC. Neither PV+ nor SST+ cell density was not affected by bedding condition or laminar region. **D-I:** cFos expression (D-E) and co-localization of cFos with PV (F-G) or SST (H-I) in cells of the layer II/III and layer V PL mPFC in NB and LB pre-adolescent offspring. Two-way ANOVA with bedding and treatment as between-subject factors were used to analyze these data. **D-E:** Cell density of cFos positive cells was significantly higher in fear-exposed animals compared to naïve controls (treatment effect: p<0.001) in both layers of the PL mPFC. **F**: Co-expression of cFos and PV in layer II/III of the PL mPFC was elevated by fear treatment (treatment effect: p<0.001), and tended to be higher in LB-exposed animals compared to NB controls (bedding effect: p=0.106). **G:**. Co-expression of cFos and PV in layer V of the PL mPFC was elevated by both fear treatment and LB exposure (treatment effect: p<0.01; bedding effect: p<0.05). **H-I:** Cell density of cFos and SST in both layers of the PL mPFC was significantly higher in fear-exposed animals compared to naïve controls (treatment effect: p<0.001), but there was no significant effect of bedding in either layer. All values are represented as mean +/- SEM. n=5-6 per group. **, p<0.01. ***, p<0.001.

For the co-expression of cFos and PV, a two-way ANOVA revealed a significant main effect of treatment (F(1,19)=19.65, p =0.0003) and a trend towards a bedding effect (F(1,19)=5.591, p =0.1066) in layer II/II of PL mPFC, with no bedding x treatment interaction (F (1, 19) = 2.349, p =0.1418) (Figure 5F). In layer V, there were significant effects of treatment (F (1, 19) = 8.405, p =0.0092) and bedding condition (F (1, 19) = 4.766, p =0.0418), without an interaction between bedding and treatment (F (1, 19) = 1.520, p =2.2326) (Figure 5G). The cFos and PV co-expression was the highest in LB offspring exposed to fear compared to NB controls and naïve animals. Co-expression of cFos and SST was significantly elevated by treatment (layer II/III: F(1,19)=21.16, p =0.0002; layer V: F (1, 19) = 29.95, p<0.0001), but was similar across bedding conditions (layer II/III: F(1,19)=0.3088 p =0.5849; layer V: F (1, 19) = 0.5431, p = 0.4701). There was no bedding x treatment interaction in either layer (Layer II/III: F(1,19)=0.4152 p =0.5271; layer V: F (1, 19) = 0.0006, p =0.9807) (Figure 5H-I).

## Discussion

In this study, we examined the effects of early life stress, in the form of the limited bedding (LB) paradigm, on the fear-induced glutamate release in the PL region of the mPFC, layer specific glutamate release probability in the mPFC and distribution of long-range glutamatergic inputs targeting the PL mPFC. In addition, we determined whether activation of local inhibitory interneurons during fear conditioning varied as a function of cortical layers in this region. Our novel findings are that LB exposure significantly enhanced fear-induced glutamatergic neurotransmission in the PL mPFC of adult males, while suppressing it in pre-adolescent males. Interestingly, these LB-induced alterations in glutamatergic transmission were not observed in female offspring of both ages. The reduced fear-induced glutamate response in pre-adolescent LB males was associated with higher activation of PV, but not SST interneurons, specifically in the layer V of the PL mPFC. At a time of maturation of PL mPFC inputs, we observed significant layer-specific effects of LB on presynaptic glutamatergic transmission, where LB exposure enhanced presynaptic glutamate release probability in the layer II/III of PL mPFC, but decreased it in the layer V. Tracing the origins of long range inputs targeting the PL mPFC, we found that LB exposure disrupted the laminar distribution of BLA and vHIP projections. These findings highlight a unique, layer-specific regulation of the corticolimbic circuit in pre-adolescent males after early stress exposure.

The mPFC receives numerous glutamatergic long-range projections from various cortical and subcortical regions, but the projections originating from the BLA and the vHIP are particularly implicated in cue-associative and contextual fear learning (Giustino & Maren 2015). Glutamate concentrations in the PL mPFC measured by *in vivo* microdialysis integrate both long range glutamatergic afferent inputs as well as local glutamatergic release. In adult males, LB exposure enhanced glutamate release in the PL mPFC during and after fear conditioning, without changes in the basal levels of glutamate concentrations, suggesting that the effects of early adversity might be observed preferentially when the corticolimbic system is challenged. Increased fear-induced PL mPFC glutamate concentrations and behavioral freezing responses in LB adult males are thought to result from hyperactivity of long-range inputs within the fear circuitry. In adults, optogenetic activation of BLA-PL mPFC projections induces freezing responses (Burgos-Robles et al 2017), while inactivation of the vHIP afferents increases PL neuronal activity to fear expression (Sotres-Bayon et al 2012). Following exposure to ELS, dendritic length and excitability of BLA pyramidal neurons is increased in adult (Eiland & Romeo 2013, Rau et al 2015), while dendritic arborization and synaptic plasticity in adult vHIP is reduced (Brunson et al 2005, Molet et al 2016), both phenomena potentially contribute to maintaining an overdrive on mPFC PL neurons and increased fear expression in adult LB male offspring.

In contrast to the adults, LB exposure tended to suppress the PL mPFC glutamate response to fear conditioning in pre-adolescent male rats. We hypothesize that this might result from LB-induced modifications in the maturation of prefrontal synaptic connections and/or from changes in the developmental trajectory of excitatory projections targeting this region. In normally reared animals, a peak in spine density in the PL mPFC is observed around PND30, together with a peak in the density of afferent projections from the BLA and the vHIP to the mPFC (Pattwell et al 2016). Research in both human and rodents suggests that exposure to early life stress accelerates the maturation of the corticolimbic circuit (Bath et al 2016, Callaghan & Tottenham 2016), potentially facilitating earlier synaptic pruning process in the mPFC of pre-adolescents. Our laboratory and others have previously demonstrated that the resting-state functional connectivity between the BLA and the mPFC is reduced in ELS-exposed pre-pubertal male offspring (Guadagno et al 2018a, Honeycutt et al 2020), possibly reflecting the advanced maturation of the BLA-mPFC reciprocal connections. Similarly, accelerated maturation of the hippocampus has also been reported in pre-weaning male mouse pups exposed to LB conditions (Bath et al 2016). Thus, it is plausible that the precocious maturation of afferent projections to mPFC in pre-adolescent LB males would advance the synaptic remodeling and the reorganization of glutamatergic afferents to the mPFC. Consistent with this hypothesis, our retrograde tracing data revealed that LB exposure induces an adult-like laminar distribution of anterior BLA projections to the PL mPFC in pre-adolescent males, with an increased proportion of neurons projecting to layer V. Although we did not observe a reduction in afferent projections to the PL mPFC in LB compared to NB pre-adolescent male offspring, previous studies reported downregulation of synaptic connectivity and plasticity in this region in early adolescents (PND35) subjected to neonatal maternal separation (Chocyk et al 2013, Majcher-Maslanka et al 2018, Monroy et al 2010). In line with these findings, we documented significant changes in presynaptic release properties of glutamatergic terminals in the PL mPFC that point to LB-induced alterations in the activity of these terminals.

As both functional characteristics of mPFC pyramidal neurons and the organization of glutamatergic afferents in this region are layer-specific (Anastasiades & Carter 2021, Song & Moyer 2018), we tested whether a layer-dependent decrease in presynaptic glutamate release in the PL mPFC could explain, at least in part, the reduced glutamate response to fear conditioning in pre-adolescent LB offspring. This was indeed the case as in layer V, where LB exposure decreased presynaptic glutamate release probability, but not in layer II/III, where LB-induced increase was observed. While the effect of ELS on PPR was drastically different between mPFC layers, this measure only cannot accurately predict overall glutamate release. The effect of ELS on glutamate responses to fear might also be dependent on layer-specific changes in distribution and activity of mPFC afferents. In pre-adolescent male rats, we found that LB exposure increased aBLA inputs to layer V of the PL mPFC, without affecting inputs to layer II/III or projections from the pBLA. For the vHIP, only NB but not LB offspring exhibited an adult-like layer patterning, suggesting that LB disrupted this normal development of laminar innervation. Although the pBLA and MDThal projections to the PL mPFC were more dense than those from the aBLA, there were no differences between bedding groups. Specific modalities and timing of ELS paradigms might differentially affect mPFC afferent development as Honeycutt et al. (2020) found that axonal arborization of BLA innervation in the mPFC of PND28 male rats was not altered by maternal separation occurring between PND2-20. While Honeycutt et al., employed anterograde tracing to assess axonal arborization of total BLA inputs, our study used retrograde tracing to quantify the number of PL mPFC projecting neurons in antero-posterior BLA subregions and thus, might reflect different maturational processes. Together, these data suggest that the anterior BLA might be more sensitive to ELS in male offspring by increasing the density of neurons projecting to the mPFC, even though axonal arborization of these projections might not yet be significantly altered at this age.

In contrast to males, fear-induced glutamate release in the PL mPFC of adult and pre-adolescent females was not significantly altered by LB conditions. Sex-dependent effects of ELS on the corticolimbic system have been well documented, with a more pronounced effect on male compared to female offspring (Bath, 2020; Walker et al. 2017), although limited data are available on the consequences of ELS on the PL mPFC in female rats. Honeycutt et al. (2020) demonstrated that maternal separation enhanced BLA-derived axonal innervation to the PL mPFC in pre- and late adolescent females, but not in males. The increased projections and stronger resting-state functional connectivity between BLA and mPFC in the ELS-exposed adolescent females might compensate for some of the detrimental effects of ELS observed in young males. In adults, estrogen may confer neuroprotective effects (Luine & Frankfurt, 2013; Page & Coutellier, 2019) against ELS-induced increases in glutamatergic transmission that we observed in adult males. The lack of a bedding effect on fear-induced glutamate secretion and freezing behavior in pre-adolescent females suggest that mechanisms conferring “resilience” to adverse conditions emerge prior to the increase in estrogen secretion at puberty. The precise nature and timing of these mechanisms needs further exploration.

Pyramidal neurons in the mPFC are heavily regulated by several types of interneurons, which can themselves be the target of long range glutamatergic projections (Nagy-Pal et al 2023, Yang et al 2021). Notably, PV and SST interneurons regulate glutamatergic afferent projections (Manz et al 2019, Urban-Ciecko & Barth 2016), and thus actively participate in the complex regulation of mPFC-mediated behaviors. Here we examined whether potential changes in inhibitory tone in the PL mPFC of LB offspring might associate with changes in glutamate concentrations during fear conditioning in pre-adolescent rats. Our results show that LB exposure increased the activation of PV, but not SST interneurons during fear conditioning, without affecting the density of these interneurons. Similarly to our results, several studies have shown that prefrontal PV interneurons might be hyperactive after chronic stress exposure in adult animals (Nawreen et al 2024; Page & Coutellier 2019). Compared to SST interneurons, the fast-spiking PV cells require high energy supplied by a high density of mitochondria, rendering this class of interneurons more vulnerable to oxidative stress in adulthood (Ruden et al 2021) and potential disruptions caused by early stress. Interestingly, we did not observe changes in PV cell density in pre-adolescent offspring exposed to LB conditions. Notably, the LB-induced increased activation of PV interneurons was significant in layer V, but not in layer II/III of the PL mPFC. It is possible that the layer-specific enhancement of GABAergic transmission in layer V might contribute to the reduced presynaptic glutamatergic transmission that we documented with our field potential recordings.

Increased intra- and extra-cortical excitatory inputs to mPFC PV interneurons have also been documented in another early stress model of post-weaning social isolation (Biro et al 2023). In our study, LB exposure enhanced activation of PV interneurons after fear conditioning in the layer V of PL mPFC. We speculate that the LB-induced increase in the density of aBLA neurons projecting to layer V of the PL area might also contribute to this enhanced PV interneuron activation.

One limitation of our electrophysiological data is that field potentials were recorded without pharmacological blockade of GABAa receptors. As a result, the layer-specific changes we observed in PPRs may involve changes in local inhibitory tone in addition to glutamate release. Early life stress has been shown to modify the function of interneurons, and our data show increased fear-induced activation of PV interneurons in layer V of the mPFC. This layer-specific enhancement of inhibitory activity might influence PPR via GABAaR and/or GABAbR mediated inhibition. Additional studies are required to identify the precise role of each receptor in this process. Together, these should allow to determine more precisely to what extent the ELS-induced changes in PPR are driven by intrinsic alterations in excitatory terminals versus local inhibitory modulation.

In conclusion, our results demonstrate that exposure to LB conditions leads to age- and sex-dependent effects on the fear-induced glutamate response in the PL mPFC. Unlike the increased glutamatergic transmission observed in adult LB males, glutamate release during fear conditioning tended to be diminished in pre-adolescent LB males. Our attempt to understand some of the mechanisms underlying these *in vivo* functional changes in pre-adolescent males revealed that LB has differential effects on PL mPFC superficial and deep layers, with respect to input density from BLA and vHIP regions, glutamate release probability and activation of PV interneurons. Thus, in this specific pre-adolescent period, LB exerts already complex effects on the structure and connectivity of the corticolimbic circuit. Additional studies at different ages and in both sexes are required to map out more precisely the various maturational processes that are affected by ELS. Although in our study glutamate responses were not modified by LB exposure in females, examining pathways that allow resilience in females is likely to enrich our understanding of the complex regulation and potential differential maturation pace of the corticolimbic circuitry in both sexes.

## Acknowledgements

The authors wish to thank Mr Luc Moquin and Dr Alain Gratton (Douglas Institute Research Center) for their help with the microdialysis experiments. The present study used the services of the Molecular and Cellular Microscopy Platform at the Douglas Institute Research Center.

