## Supplemental Figure 1-1 for "Layer-specific glutamatergic inputs and Parvalbumin interneurons modulate early life stress induced alterations in prefrontal glutamate release during fear conditioning in pre-adolescent rats"

**Figure 1-1:** Freezing behavior during fear conditioning intervals in NB and LB offspring.

| Freezing behavior (%) |  | **Adult Male*** |  |  | Adult Female |  |
| --- | --- | --- | --- | --- | --- | --- |
|  |  | NB | LB |  | NB | LB |
| Interval 1 |  | 43.51±6.15 | 50.51±6.97 |  | 38.82±9.47 | 34.13±6.17 |
| Interval 2 |  | 65.92±7.35 | 74.17±6.04 |  | 56.68±10.11 | 64.42±6.36 |
| Interval 3 |  | 79.61±5.27 | 81.20±3.69 |  | 60.42±7.12 | 72.92±4.89 |
| Interval 4 |  | 44.28±8.26 | 76.43±7.39 |  | 66.24±11.07 | 56.46±5.88 |
| Interval 5 |  | 69.71±7.54 | 75.06±7.15 |  | 58.63±5.84 | 62.16±5.13 |
| Interval 6 |  | 70.96±5.17 | 76.71±7.64 |  | 43.88±6.77 | 51.81±5.87 |
| Interval 7 |  | 45.10±10.06 | 69.93±7.27 |  | 62.87±7.93 | 44.51±7.81 |
| Interval 8 |  | 68.28±5.51 | 79.62±4.51 |  | 56.77±6.37 | 69.57±5.74 |
| Interval 9 |  | 54.56±8.19 | 66.50±7.68 |  | 40.58±6.35 | 59.63±6.96 |
| Interval 10 |  | 40.36±8.32 | 64.08±6.09 |  | 43.06±8.89 | 43.02±5.14 |
|  |  | Pre-adolescent Male |  |  | Pre-adolescent Female |  |
|  |  | NB | LB |  | NB | LB |
| Interval 1 |  | 42.29±6.82 | 36.70±6.09 |  | 32.86±4.45 | 35.74±9.43 |
| Interval 2 |  | 50.19±9.34 | 56.19±6.33 |  | 62.22±10.51 | 63.22±5.53 |
| Interval 3 |  | 63.35±7.54 | 72.01±6.79 |  | 70.26±10.58 | 69.12±7.85 |
| Interval 4 |  | 50.90±10.18 | 58.19±8.67 |  | 64.32±9.88 | 62.07±9.92 |
| Interval 5 |  | 65.64±7.69 | 67.65±8.38 |  | 62.53±9.50 | 67.25±7.30 |
| Interval 6 |  | 69.07±7.01 | 71.58±8.78 |  | 74.62±7.31 | 76.05±5.28 |
| Interval 7 |  | 63.49±9.25 | 61.69±8.75 |  | 52.55±10.32 | 60.77±9.84 |
| Interval 8 |  | 77.15±5.19 | 73.62±6.57 |  | 71.40±5.00 | 74.63±5.67 |
| Interval 9 |  | 57.33±7.44 | 59.62±10.26 |  | 41.90±8.69 | 61.19±10.06 |
| Interval 10 |  | 54.69±7.69 | 60.18±8.41 |  | 60.13±7.30 | 56.24±8.74 |
| * p < 0.05 NB vs LB |  | |  |  | |  |

Percentage of freezing time across ten tone-shock intervals during in vivo microdialysis in adult male, adult female, pre-adolescent male, and pre-adolescent female offspring. Two-way ANOVAs were conducted with bedding condition as a between-subject factor and interval as a within-subject factor. Fear conditioning significantly increased freezing behavior over ten intervals in all groups of animals (interval effects: p<0.001). A significant main effect of bedding was only observed in adult males (F(1,25)=6.377, p=0.018), where LB animals displayed significantly more freezing compared to NB controls. No significant interaction between bedding x interval was observed. All values are represented as mean +/- SEM. n=9-17 animals per group. *, p<0.05.
