## Supplementary figures and images for "Layer-specific glutamatergic inputs and Parvalbumin interneurons modulate early life stress induced alterations in prefrontal glutamate release during fear conditioning in pre-adolescent rats"

### Supplemental Figure 4-1

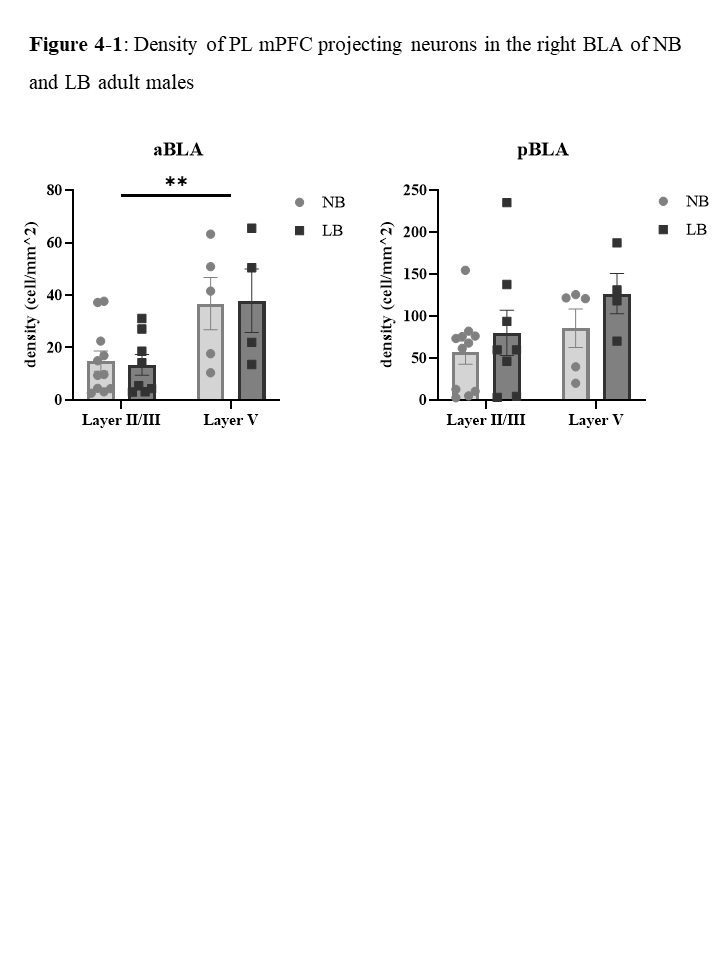
